# Allosteric Site Prediction Using Protein Language Models and Orthosteric Conditioning

**DOI:** 10.1101/2025.06.27.662060

**Authors:** R. C. Eccleston, N. Furnham

## Abstract

Allosteric modulators as therapeutics offer many advantages over orthosteric modulators, including improved selectivity and tunability. However, identifying and characterising allosteric sites remains a major challenge both experimentally and computationally. Accurate prediction of allosteric binding sites is critical to facilitate allosteric drug discovery. Here, we evaluate three strategies to predict allosteric sites using pre-trained protein language models (pLMs), Ankh, ProtT5 and ESM-2, which are trained on sequence information alone. First, a classifier was trained using static embeddings extracted from each pLM. Second, parameter-efficient fine-tuning was implemented using LoRA with focal loss to account for the sparse positive labels and lack of data. Finally, a structure-aware conditioning mechanism was introduced, whereby orthosteric binding sites are encoded and integrated directly into the input embeddings to enable the model to predict allosteric binding sites, conditioned on the knowledge of the orthosteric binding pocket. This approach, which captures functional dependencies between the othosteric and allosteric binding sites, improves upon the performance of the first two methods and achieves results comparable to the leading structure-based allosteric site predictors.

## Introduction

Allostery is critical to the regulation of protein activity and occurs when the binding of an effector modulator at site distal from the active (or orthosteric) site, alters protein conformation and dynamics [1]. Proteins exist as dynamic conformational ensembles fluctuating between multiple structural states. The native state is usually the most stable and thus most populated under physiological conditions and is often the conformation associated with the protein’s primary function. However, proteins can also transiently adopt alternative conformations with different functional properties and lower stability. The binding of an allosteric modulator, shifts the equilibrium of the conformational ensemble by stabilising alternative states, either enhancing (positive allosteric modulation) or inhibiting (negative allosteric modulation) the protein’s activity [1, 2, 3].

Rational drug design mainly focuses on developing drugs that bind at the active site and prevent the substrate or cofactor from binding. However, for many target proteins, the active site is considered undruggable, often because the active site is highly conserved across protein families, leading to undesirable off-target effects. Allosteric sites, however, are less conserved across protein families and so allosteric drugs can be more selective and thus more tolerated with fewer side effects [4]. Furthermore, whereas orthosteric drugs typically act as full-agonists/antagonists, turning protein activity on/off, allosteric drugs can be used to fine-tune modulation by either enhancing or inhibiting activity in the presence of the endogenous ligand, allowing natural regulation to be preserved. For example, benzodiazepines act as positive allosteric modulator as they enhance the effect of GABA_A_, a neurotransmitter that inhibits neuronal activity [5].

The therapeutic potential of allosteric modulators has driven growing interest in developing drugs that target allosteric binding sites, leading to the approval of several allosteric drugs – catalogued in resources such as the Allosteric Site Database (ASD) [6]. However, identifying and developing allosteric drugs remains challenging because allosteric sites and their functional roles are often unknown.

Allosteric binding sites are typically discovered serendipitously, as they are difficult to study experimentally. To address this, computational methods have been developed to aid in allosteric site identification [7], yet predicting allosteric pockets remains substantially more difficult than orthosteric ones. Allosteric sites tend to be less evolutionarily conserved, involve fewer residues and exhibit more ambiguous, non-pocket-like geometries, limiting the effectiveness of traditional approaches based on sequence conservation, co-evolution or structural pocket detection tools, like FPocket [8]. Allosteric sites are usually closer to the surface making them harder to distinguish from other random surface grooves or shallow depressions, increasing the risk of false positives in prediction. Moreover, allosteric sites often only become apparent upon ligand binding, meaning high-resolution structures revealing these pockets are frequently unavailable. Therefore, unlike orthosteric sites, few allosteric sites are fully annotated, or experimentally validated making supervised learning harder.

A variety of supervised machine learning (ML) models have been developed to predict allosteric binding sites using structure-based features, such as physiochemical properties, Normal Mode Analysis (NMA) and network representations (see [7] for a review). Resources like the ASD and ASBench [6, 9, 10, 11, 12] provide curated datasets of protein-modulator complexes, including experimentally determined protein structures and annotated allosteric binding sites which enable the construction of labelled training data. Most existing models focus on predicting the binding pocket - for example, Allosite [13], AlloPred [14], AlloSitePro [15], ALLO [16], PASSer [17], PASSer2.0 [18], PASSerRank [19], AlloReverse [20]. Others aim to identify the specific allosteric residues involved, such as NACEN [21] and AR-Pred [22], or broader structural regions where allosteric modulation might occur, as in TopoAlloSite [23] which predicts protein domains or interfaces. Where evaluation metrics are provided to enable comparison, PASSer appears to be the best performing model to date with accuracy 0.97 and recall 0.85 [7].

All existing allosteric prediction models to date use structure-based features that rely on detailed geometric and physiochemical descriptors derived from high-resolution 3D structures. However, our approach uses pretrained protein language models (pLMs), exclusively trained on large datasets of diverse protein sequences. pLMs have demonstrated the ability to infer structural and functional properties directly from sequence, such as amino acid contacts, secondary structure [24], protein folding [25], post-translational modifications [26], and biophysical properties such as solubility and fluorescence, and have also shown promise in predicting the effects of protein variants [26, 25]. pLMs were inspired by the success of large language models (LLMs) in natural language processing [27]. Built on the attention-based Transformer architecture [28], models like BERT (Bidirectional Encoder Representations from Transformers) [29], trained on massive text datasets, learn context-aware representations of sequence data, and the same architecture can be adapted to train on protein sequences. In structural biology, the sequence-structure-function paradigm posits that the sequence of amino acids in a protein determines its 3D structure and function. Analogous to human language, where meaning arises from the order and context of words, protein sequences exhibit grammar-like properties, where conserved motifs and residue co-occurrence encode biological meaning [30]. Therefore, pLMs trained on billions of protein sequences can learn latent representations that implicitly capture structural and functional constrains – without requiring explicit structural input.

We investigate if pre-trained pLMs - ProtT5 XL [26], Ankh Large [24] and ESM-2 3B [25] - can identify allosteric binding sites. This task is particularly challenging due to the limited size of the training dataset and a severe class imbalance, where positive examples (allosteric residues) comprise only ∼6% of the total labels. To address this, we explore three strategies:

First, we extract token-level embeddings from the final hidden layer of each pLM (i.e. the vector representation of each amino acid in a protein sequence) and train a classifier to identify allosteric residues based on these embeddings. In theory, if the embeddings adequately capture sequence-level relationships, the classifier should be able to learn discriminative features for allosteric site prediction. To improve performance, we also experiment with transfer learning by pre-training the classification head on the PDBbind dataset [31, 32] - a large dataset of ligand binding sites - before training the classifier on the ASD dataset. However, both direct classification and transfer learning variants result in poor performance across all three pLMs, falling short of existing structure-based allosteric site predictors.

Second, we performed parameter-efficient fine-tuning using Low Rank Adaptation (LoRA) whereby most of the model’s parameters were frozen, and only a small fraction of the weights were updated. This enables efficient training on limited resources. To optimise performance, we performed hyperparameter tuning on the LoRA layers and other parameters such as weight-decay and compared weighted cross-entropy loss and focal loss to determine which is best at handling severe class imbalance. Fine-tuning leads to notable improvements over the first method, but performance still falls short of state-of-the-art tools.

Third, we introduce a structure-aware conditioning mechanism, leveraging a subset of the ASD dataset that includes both allosteric and orthosteric binding site annotations. We encode orthosteric site information and inject it directly into the input embeddings, allowing the model to condition its allosteric site prediction on the knowledge of the orthosteric binding pocket. This approach is biologically motivated: allosteric modulation requires an orthosteric site to exert functional effects i.e. there would be no allosteric site without an orthosteric site. By incorporating this dependency, the model effectively learns a functional coupling between the two sites and yields the best results of the three methods presented here, with performance comparable to the leading structure-based allosteric site predictors.

## Results and Discussion

### Method 1: Classification Using Raw Token-Level Embeddings from Pretrained pLMs

We evaluated three pre-trained pLMs – Ankh large, ProtT5 XL and ESM-2 3B – for their ability to predict allosteric binding sites at the residue-level. Briefly, for each pLM, we extracted the final hidden state token-level embeddings (without fine-tuning the model weights) for every protein sequence in the training set. These embeddings were then used as input features to train a supervised classifier to predict allosteric binding residues from labelled training data (Figure 1a). We used the `ConvBertForTokenClassification` model provided by the Ankh repository as the classifier, trained with class-weighted cross-entropy loss to address label imbalance. This classifier combines convolutional layers with transformer-based multi-head self-attention, enabling it to learn both local residue-level features and long-range contextual dependencies from the embeddings.

**Figure 1.**
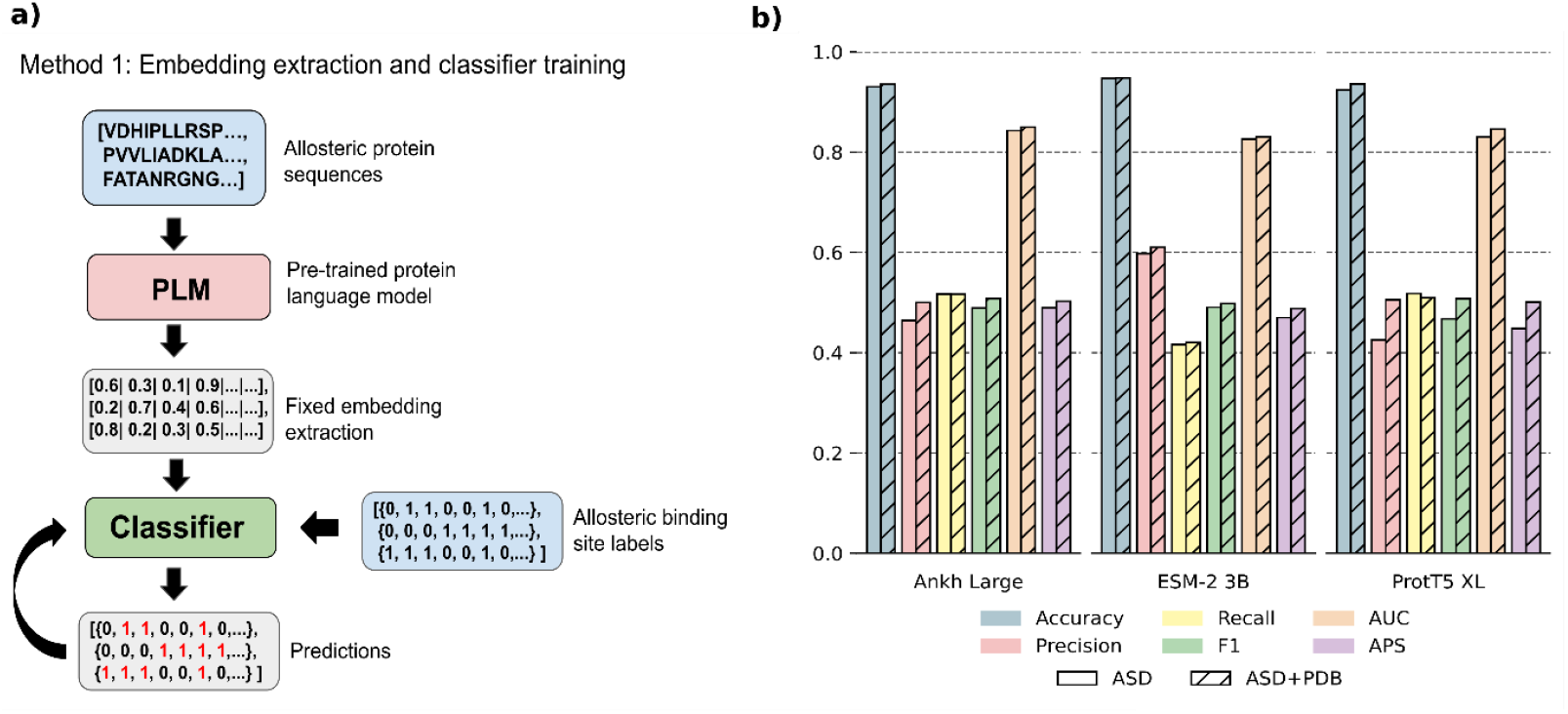
a) Schematic of pipeline for Method 1, showing input of allosteric protein sequences to pLM, fixed embedding extraction, training of classifier with fixed embeddings and residue level classes and predictions using trained classifier, b) performance of Ankh Large, ESM-2 3B and ProtT5 XL predicting allosteric binding sites using Method 1, bars are coloured by metric and shaded by dataset. Hatched bars indicate transfer learning was used to pre-train classifier on the PDB bind dataset.

**Figure 2.**
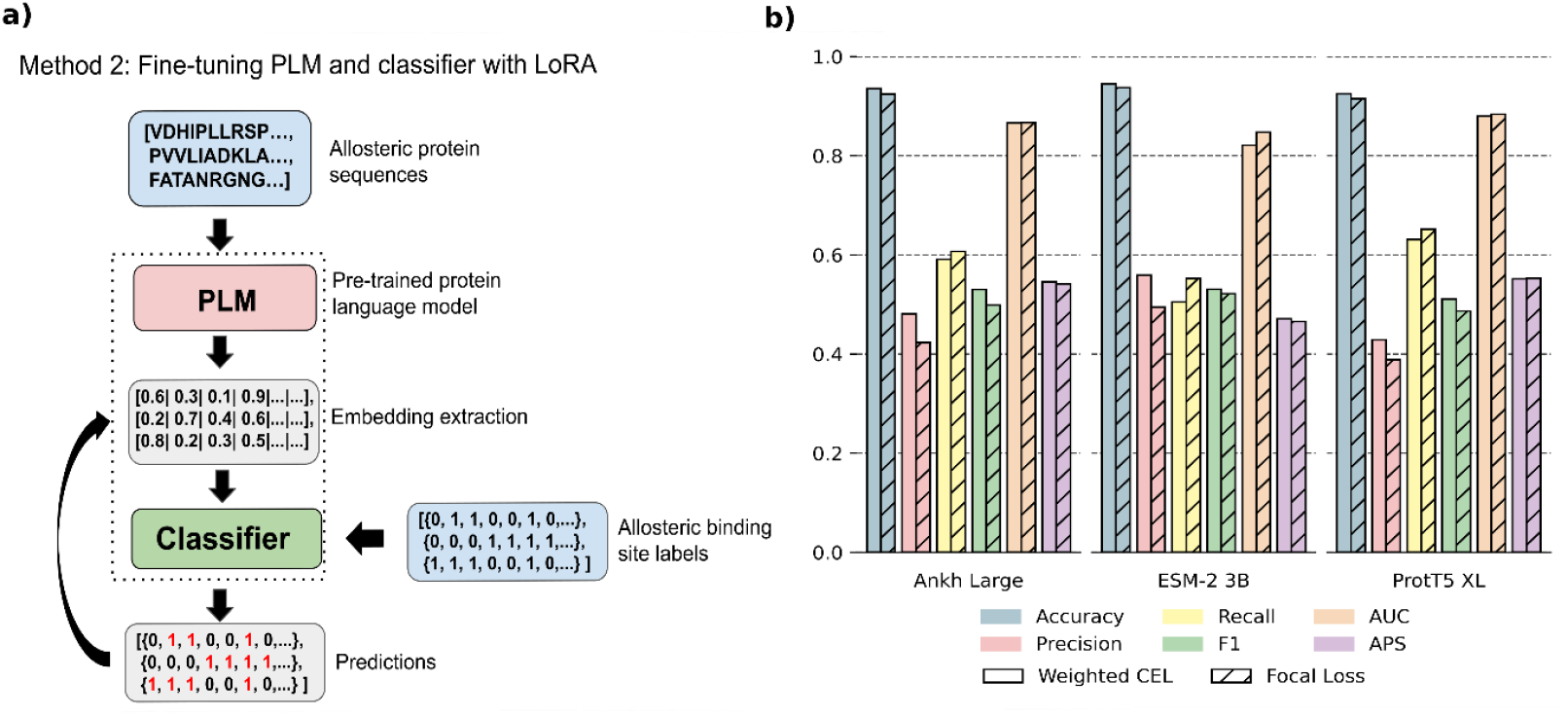
a) Schematic of pipeline for Method 2, showing fine-tuning of both PLM and classifier weights, b) results of fine-tuning Ankh Large, ESM-2 3B and ProtT5 XL. Bars are coloured by metric and shaded by loss type. Hatched bars indicate focal loss, solid bars indicate weighted cross-entropy loss.

We trained the classifier using data from the Allosteric Site Database (ASD) and split the data into training, validation and test sets. To reduce data leakage and ensure the model is generalisable, we engineered the train/validation/test split so that none of the sequences in the training dataset had a greater than 30% sequence identity with sequences in the validation and test set.

Separately, we also used transfer learning to first train the classification head on ligand binding site data from PDB Bind before then training on the ASD dataset to determine if this resulted in improved performance. We chose to pre-train the classifier on data from PDBbind because it was previously reported to improve prediction of cryptic binding sites (an allosteric site that is hidden in the protein’s native structure, which is then revealed when a ligand binds at a different site and causes conformational changes) [33]. However, cryptic binding sites tend to be as evolutionarily conserved as orthosteric binding sites and are structurally deep and enclosed like orthosteric sites [34], so whilst they are mechanistically allosteric, their pocket geometry and structure may be more similar to orthosteric pockets on formation. We reasoned that pre-training the classifier weights on any form of additional binding data may improve performance when training on the smaller ASD dataset (after pre-processing the PDBbind dataset contained around 14,000 binding sites compared to around 3,000 for the ASD dataset). Hyperparameter values for Method 1 can be found in Figure S1.

To compare the performance of the pLMs, we calculated several evaluation metrics: accuracy, precision, recall, F1, AUC-ROC and average precision score (APS) (Figure 1b). ESM-2 3B has the highest precision and accuracy, with and without transfer learning, but the lowest AUC-ROC, APS, F1 and recall. Ankh large has the highest AUC-ROC, APS, F1 and recall, whilst ProtT5 XL fell in the middle of the other two models for most metrics, except APS where it was lowest.

Both accuracy and AUC-ROC values were high for all models, with small improvements in these metrics observed with transfer learning. Accuracy ranged from 0.923 to 0.947 for ASD only and 0.935 to 0.948 with transfer learning. Similarly, AUC-ROC values were between 0.826 to 0.843 for just ASD data and 0.830 to 0.850 for transfer learning. Although accuracy and AUC-ROC are high for all three pLMs, these metrics can be misleading in the context of significant class imbalance. In our dataset, the positive class – residues labelled as allosteric – constitutes only around 6% of all residues. As a result, a model could achieve high accuracy simply by predicting all residues as non-allosteric (i.e., all 0s). Similarly, AUC-ROC may be high (close to 1), if the model ranks positive examples higher than negative ones overall, even if classification performance at a specific threshold is poor.

All three of the threshold-dependent metrics – precision, recall and F1 – indicate low-to-middle performance at classifying allosteric sites. Recall — the proportion of actual positives correctly identified — remained relatively unchanged with transfer learning, ranging from 0.416 to 0.518 for models trained only on ASD, and from 0.423 to 0.517 after transfer learning, using a classification threshold of 0.5. This suggests that even the best-performing model, ProtT5 XL, identifies just over 50% of the true allosteric residues.

In contrast, transfer learning did improve precision — the proportion of predicted positives that are correct — which increased from 0.426–0.598 (ASD only) to 0.501–0.610 (after transfer learning). This means that in the best case, 61% of residues predicted as allosteric were indeed true positives, suggesting that transfer learning helps reduce the number of false positives without compromising recall.

The improvement in precision resulted in an improvement in F1 – the harmonic mean of the precision and recall – with values of 0.467 to 0.491 for ASD only and 0.498 to 0.508 with transfer learning, at a threshold of 0.5. This again indicates the model is correctly predicting about half of the allosteric sites, with relatively high rates of false positives and false negatives.

The values of precision, recall and F1 will change depending on the classification threshold. Therefore, the average precision score (APS) may be the best metric to directly assess model performance in the context of severe class imbalance as it offers a threshold-free summary of the trade-off between precision and recall. APS is better suited for imbalanced classification tasks than other threshold-free metrics such as accuracy or ROC-AUC, as it reflects how effectively the model ranks true allosteric residues among the majority negative background. The highest APS (0.502) was observed for Ankh Large trained with transfer learning, whilst the lowest was ProtT5 XL trained without transfer learning, with an APS value of 0.448.

Without transfer learning, APS values ranged from 0.448 to 0.490 across all three pLMs. Applying transfer learning resulted in a modest improvement, with APS values increasing to between 0.488 and 0.502. These values indicate that, on average, fewer than half of the predicted allosteric sites are correct — that is, the number of true positives is roughly comparable to the number of false positives.

This suggests the raw token embeddings from the pLMs do not sufficiently capture the features needed to distinguish allosteric sites, and that the classifier trained on them has little predictive power.

### Method 2: Parameter-Efficient Fine-Tuning of pLMs with LoRA

To enhance the predictive power of the pLM embeddings and address both the limited availability of allosteric site data and the significant class imbalance, we fine-tuned each pLM with the `ConvBertForTokenClassification` head as a classifier. We performed hyperparameter turning on the LoRA layer parameters (rank - r, alpha - α), the learning rate and weight decay and experimented with focal loss to better address large class imbalance. The hyperparameter values for this method can be found in Figures S2-S6.

Accuracy remained largely unchanged compared to Method 1, with values that ranged from 0.924 to 0.945 with weighted cross-entropy loss and from 0.915 to 0.937 with focal loss. There was a small increase in AUC-ROC values compared to Method 1, which ranged from 0.82 to 0.878 with weighted cross-entropy loss and from 0.847 to 0.884 for focal loss.

Recall increased for all models compared to Method 1, with values of 0.505 to 0.631 for weighted cross-entropy loss and 0.552 to 0.651 for focal loss. At a classification threshold of 0.5, the recall this suggests the best fine-tuned model can correctly identify up to 65% of allosteric sites in the dataset. For both losses, ProtT5 XL had the highest values of recall and ESM-2 3B had the lowest. The increase in recall resulted in a drop in precision compared to Method 1, with values between 0.429 to 0.481 for cross-entropy loss and 0.389 to 0.466 for focal loss, with ESM-2 3B having the highest precision across both losses and ProtT5 XL having the lowest. Overall, F1 improved compared to Method 1, ranging from 0.511 to 0.532 for weighted cross-entropy loss and from 0.487 to 0.522 for focal loss, with Ankh Large having the highest F1 value overall.

In terms of our representative summary metric APS, ProtT5 XL performed best across both loss functions, with the highest of APS of 0.553 achieved using focal loss compared to 0.552 for weighted cross-entropy loss. Across all models, APS values ranged from 0.471 to 0.552 for weighted cross-entropy loss and 0.466 to 0.553 for focal loss, with ESM-2 3B (trained with identical hyperparameters as ProtT5 XL) having the lowest APS values for both losses. There is an almost 11% increase in APS compared to Method 1 and on average, the best performing fine-tuned model correctly predicts allosteric sites over 55% of the time.

When comparing the performance (accuracy and recall) of our best fine-tuned model against existing allosteric site predictors, we found it ranked 4th in accuracy and 6^th^ in recall (Table 1), outperforming several models that rely on both structural and sequence-based features.

**Table 1.** Comparison of performance our orthosteric-conditioned model (Method 3) with that of existing allosteric prediction models.

| MODEL | YEAR | OUTPUT | ACCURACY | RECALL |
| --- | --- | --- | --- | --- |
| ALLOSITE | 2013 | Pockets | >95 | >83 |
| ALLOPRED | 2015 | Pockets |  |  |
| ALLOSITEPRO | 2017 | Pockets |  | 52 |
| ALLO | 2018 | Pockets |  |  |
| NACEN | 2018 | Residues | 83 | 30 |
| AR-PRED | 2019 | Residues | ~70 | ~80 |
| PASSER | 2021 | Pockets | **97.4** | **84.7** |
| PASSER2.0 | 2022 | Pockets |  | 61.6 |
| PASSERRANK | 2023 | Pockets | 96.8 | 66.2 |
| ALLOREVERSE | 2023 | Pockets |  | 71 |
| TOPOALLOSITE | 2023 | Location | 76 |  |
| METHOD 3: ESM-2 3B | 2025 | Residues | 95.3 | 78.8 |
| METHOD 3: ANKH-LARGE | 2025 | Residues | 92.4 | 83.6 |

These improvements across key metrics demonstrates fine-tuning with LoRA substantially enhances with informativeness of the PLM embeddings for this task when compared to the raw pretrained embeddings

### Method 3: Conditioning pLMs with Orthosteric Sites Embeddings as an Additional Feature

Predicting allosteric sites using pLMs is inherently challenging, as these models are trained solely on sequence data and lack structural or dynamic context. This is a key limitation, since allosteric regulation often involves shifts in the conformational ensemble and structural coupling between the allosteric and orthosteric binding sites.

To address this, we introduced a structure-aware conditioning mechanism that incorporates orthosteric site information into the model input (Figure 3a). Specifically, we leveraged a subset of the Allosteric Database (ASD), which provides both allosteric and their corresponding orthosteric binding sites. As with previous methods, we generated token-level allosteric site labels, and additionally derived orthosteric site masks — binary vectors (0/1) marking residues in the orthosteric pocket.

**Figure 3.**
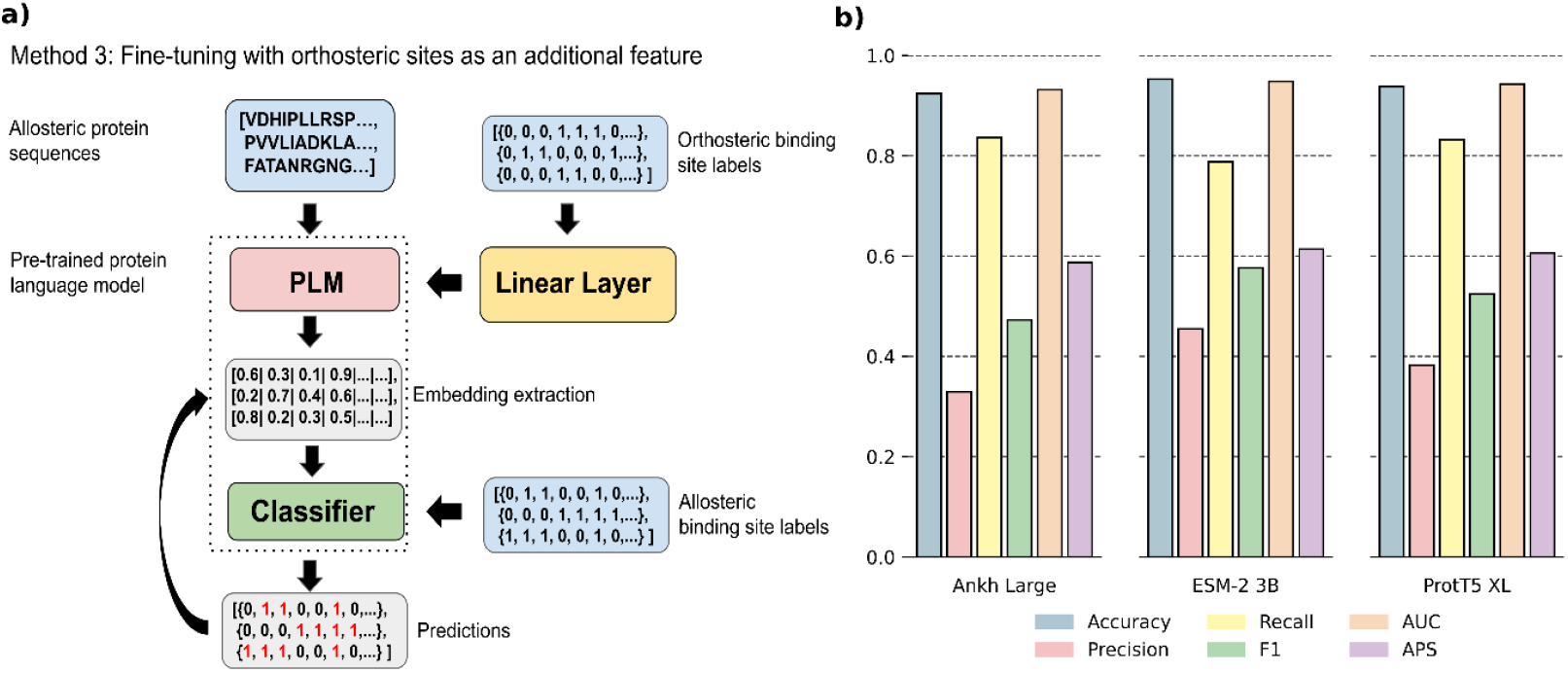
a) Schematic of pipeline for Method 3, showing additional of orthosteric binding site embeddings to condition allosteric site prediction. b) Results of orthosteric-conditioned model for the three pLMs. Bars are coloured by metric.

During fine-tuning, each orthosteric mask was linearly projected to match the embedding dimensionality and added to the input token embeddings. This conditioned the model’s representation of each residue on the spatial proximity and identity of the orthosteric pocket, allowing the model to learn functional dependencies between orthosteric and allosteric sites. We chose to implement this conditioning mechanism over multi-task learning due to the limited number of sequences in the joint allosteric-orthosteric dataset (around 1,900 sequences) which we hypothesized might lead to unstable training or insufficient learning signal for either task. Embedding conditioning allowed us to incorporate orthosteric site information without introducing an additional loss term, reducing the risk of overfitting or task interference. The hyperparameter values for this method can be found in Figures S7-S9.

Because the orthosteric-labeled dataset contains approximately 1,900 proteins — roughly half the size of the dataset used for Methods 1 and 2 — we initialized this model with the best-performing model from Method 2. This transfer-learning strategy allowed the model to benefit from prior training on the larger dataset before honing its predictions with orthosteric conditioning. This method suffered from the potential to rapidly overfit to the training data, and so to combat this, we experimented with increasing dropout and weight decay and reduced the number of training epochs.

Empirically, this method achieved the strongest recall (0.766-0.836) and the best APS (0.587-0.614) of all approaches, demonstrating improved sensitivity to the true allosteric sites and better overall ranking of positive residues (Figure 3b). However, this gain in sensitivity came at the cost of precision, which dropped to 0.329-0.455, while F1 improved, ranging from 0.473-0.577. Accuracy increased slightly for all pLMs with a range of 0.924-0.953 and there was a significant improvement in AUC-ROC with a value of 0.932-0.948, suggesting an improvement in the model’s ability to distinguish allosteric residues from non-allosteric residues across all classification thresholds.

While the embeddings from the orthosteric-conditioned model are more informative for detecting true allosteric sites—evidenced by the substantial increase in recall and AUC-ROC —they also result in more false positives, lowering precision. This suggests the model is more generalisable, and that embeddings conditioned on orthosteric site information are more informative for predicting allosteric sites, likely because they capture functional and spatial dependencies that are not available from sequence alone. However, this improvement comes at the cost of overpredicting allosteric sites. Consequently, this model is best suited for use cases prioritizing sensitivity, where identifying all potential allosteric sites is important and downstream filtering can be applied to improve precision.

ESM-2 3B was performed best in terms of accuracy, precision, AUC-ROC, F1, and APS, whilst Ankh-large had the best recall. When benchmarked against existing allosteric site predictors, we used the performance table compiled in [7], which reports the accuracy and recall each model (Table 1). Our best model in terms of accuracy was ESM-2 3B, ranked 3^rd^ in accuracy and 4^th^ in recall. Our best model in terms of recall was Ankh-large, which ranked 4^th^ in accuracy and 2^nd^ in recall. These results demonstrate fine-tuning with orthosteric conditioning outperforms the majority of existing models that rely on structural, dynamic and sequence-based features.

## Conclusions

This study evaluated three distinct approaches for predicting allosteric binding sites using protein language models (pLMs). Our results demonstrate that fine-tuning PLMs with parameter efficient methods, such as LoRA, results in embeddings that are more informative compared to using raw pretrained embeddings alone. We also found our novel structure-aware conditioning mechanism in which orthosteric site information is incorporated as an additional feature further enhanced the model’s sensitivity, achieving the highest recall and average precision score among all methods test here.

Despite these advances, predicting allosteric sites remains inherently difficult in general, especially compared to orthosteric sites, as they are less evolutionary conserved, the pockets involve fewer residues and are less well-defined structurally. Furthermore, unlike orthosteric sites, few allosteric sites are fully annotated, or experimentally validated, making supervised learning harder. It also presents a particularly difficult task for pLMs specifically, as they tend to learn local and sequential patterns, not the global dynamics that govern allostery. This challenge likely explains the relatively poor performance of Method 1, where classifiers trained on raw PLM embeddings struggled to distinguish allosteric residues from the vast majority of non-allosteric residues. The low precision and recall in Method 1 emphasises the need for additional context or fine-tuning to extract meaningful embeddings.

Our orthosteric-conditioned model showed promising gains in recall, suggesting that integrating spatial and functional dependencies between orthosteric and allosteric sites enables pLMs to better learn the characteristics of allosteric residues, which is more difficult to do with sequence-based information alone, due to the inherently dynamic nature of allosteric modulation. However, this improvement came at the expense of precision, with increased false positives indicating a trade-off between sensitivity and specificity. This suggests the model is well-suited for applications prioritising the identification of all possible allosteric sites but further downstream refinement might be necessary for tasks requiring high precision. Moreover, this model is restricted to use cases where the orthosteric site is known or can be confidently predicted.

The limited size and class imbalance of the available allosteric datasets pose constraints on model training and evaluation, likely reducing generalisability. Future work could expand on our orthosteric-conditioning model and embedding pLMs with dynamic structural data, such as normal modes, flexibility and energetic couplings, which are not easily inferred from sequence alone but would further enhance predictive power. Alternatively, embeddings extracted from pLMs trained to be maximally informative for allosteric site prediction, such as the models presented here, could be used as features to a larger predictive model combining sequence and structural information to maximise predictive accuracy. Additionally, as more diverse allosteric datasets become available, models using sequence information alone will become more powerful, possibly overcoming the current limitations due to data availability constraints.

Overall, this work illustrates the potential of combining PLMs with relevant structural conditioning and efficient fine-tuning techniques to improve allosteric site prediction. Our findings provide a foundation for developing more sensitive and practical computational tools to aid drug discovery and deepen understanding of allosteric regulation.

## Methods

### Pre-trained pLMs

We used three state-of-the-art pLMs in this study: Ankh-large, ESM2 3B and Prot T5 XL. Ankh-large and Prot T5 XL are based on the T5 (Text-to-Text Transfer Transformer [35]) encoder-only architecture. The T5 encoder consists of a stack of transformer blocks with multi-headed self-attention layers, feedforward layers, layer normalisation and residual connections. The encoder is trained using a denoising span-masking objective, where lengths of tokens are masked so the model can learn to reconstruct them and learn rich contextual embeddings. Prot T5 XL is from Rostlab and uses a large 3B parameter model, trained on the UniRef50 dataset. Ankh-large uses a smaller encoder (1.5B parameters) but was trained on a large, diverse datasets spanning more than 60B residues in total to improve generalisability. ESM-2 (3B parameter model) was developed by Meta AI and follows standard BERT-style transformer encoder-only architecture [28], with 3B parameters and 36 layers, trained using masked language modelling (MLM) on millions of sequences from UniProt.

### Datasets

For the allosteric data we used the most recent ASD Release (ASD_Release_202306_XF, found here: https://mdl.shsmu.edu.cn/ASD/module/download/download.jsp?tabIndex=1), as the allosteric data in Methods 1 and 2. This data is provided as a set of html files, one for each protein in the dataset, in which information on the protein, the allosteric modulator and the allosteric binding sites are given. The allosteric modulators can be ligands, peptides, ions as well as protein-protein interactions.

Some of the entries were missing information on the allosteric sites. Therefore, to make the dataset more complete and obtain allosteric binding sites when they are not given in the dataset, where possible we extracted the allosteric binding sites from the allosteric pdb files (when these were provided in the html files), under the assumption that any residue within 7A of a ligand atom was an allosteric binding site.

We used the 2020 refined version of the PDBbind Database (found here: https://www.pdbbind-plus.org.cn/) as the ligand binding data used for transfer learning in Method 1. This dataset provides the structures of proteins and their ligands separately, along with a pdb file containing just the pocket residues involved in the ligand binding).

We also extracted the orthosteric-allosteric site subset data from the table from the ASD (found here: https://mdl.shsmu.edu.cn/ASD/module/site/site.jsp). This dataset provides allosteric binding sites annotated with their orthosteric binding sites where that information exists. This data was used to create orthosteric labels in the same way as above for use in the orthosteric conditioning model.

For all datasets, amino-acid level labels created for each protein sequence where each amino acid is represented by either a 0 or a 1 where 1 represents a binding site (either allosteric or orthosteric) and 0 represents non-binding site.

We performed a train/validation/test split on each dataset (0.8/0.1/0.1), using MMSeq [36] to ensure sequences in the test and validation set had >30% sequence identity with the sequences in the training dataset. The training and validation set were used during model development, while the test set was held out for final evaluation to assess generalisability (Tables 2-4).

**Table 2.** Size of ASD datasets used for all models in Method 1 and Method 2.

| *Dataset* | *Size* |
| --- | --- |
| *train* | 3206 |
| *validation* | 247 |
| *test* | 246 |

**Table 3.** Size of PDBbind datasets used for all models when transfer learning in Method 1.

| *Dataset* | *Size* |
| --- | --- |
| *train* | 12145 |
| *validation* | 989 |
| *test* | 989 |

**Table 4.** Size of orthosterc-allosteric datasets used for all models when training with orthosteric conditioning in Method 3.

| *Dataset* | *Size* |
| --- | --- |
| *train* | 1122 |
| *validation* | 142 |
| *test* | 142 |

### Model Training

For all methods, we loaded each model from publicly available pre-trained checkpoints on Hugging Face.

For Method 1, we generated embeddings for protein sequences in both the ASD and PDBbind datasets by passing the tokenized amino acid sequences through the pLM and extracting the final hidden layer representations. These embeddings were then used as input to the classifier, along with the residue-level label lists, to predict allosteric sites. When applying transfer learning, the classifier trained with the PDBbind data was saved and used as the starting point for the training with the ASD data. We performed limited hyperparameter tuning on the learning rate and weight decay; once selected, these values were held constant across runs for all three pLMs.

For Method 2, we fine-tuned the pLMs directly using Low-Rank Adaptation (LoRA). We performed limited hyperparameter tuning on LoRA parameters (rank and alpha) for each pLM and experimented minimally with adapter placement (e.g. in query, key, value and dense layers). Prot T5-XL and ESM2 3B were fine-tuned using mixed precision, whereas Ankh-large only supported full precision.

For Method 3, incorporated information about the orthosteric site locations by conditioning the model on orthosteric site labels. These binary labels were first passed through a linear layer to transform them into dense vectors with the same dimensionality as the model’s token-level embeddings. The resulting orthosteric embeddings were then added element-wise to the original input embeddings prior to the forward pass through the model. This conditioning mechanism allows the model to leverage prior knowledge of the orthosteric site locations when predicting allosteric residues, potentially enabling it to capture spatial or functional dependencies between the two types of sites.

### Transfer learning

We used transfer learning to try to improve the performance of the model. In Transfer Learning, a model is pre-trained using data from a different but related task, in the hope that it will improve the performance of the model when trained for the target task. Transfer learning is usually used in cases where the dataset used for the target task is small or limited in some way. Here, we used transfer learning to pre-train the classifier weights in Method 1 on ligand binding sites from the PDBbind dataset in the hope that it would improve performance when trained on the smaller ASD dataset.

### Loss Functions

To address the substantial class imbalance in our dataset, we compared the performance of two loss functions commonly used in situations of large class imbalances in the training data: weighted cross-entropy loss and focal loss.

Weighted cross-entropy loss is a modified form of standard cross-entropy loss that assigns higher weights to under-represented classes, encouraging the model to pay more attention to the minority class during training. The class weights were computed directly from the class distribution in the training data.

Focal loss extends the standard cross-entropy loss by introducing a modulating factor to amplify the impact of incorrectly predicted samples whilst reducing the loss impact for correctly predicted samples. This forces the model to concentrate on learning harder to learn samples and so is useful in cases of large class imbalances where there is a large disparity in the frequency of the majority and minority

## Code and Data Availability

All code and data used in this project can be found in the following repository: https://github.com/RCEccleston/PLM_Allosteric_Classification

## Supplementary Hyperparameter values

### Method 1

**Table S1.**
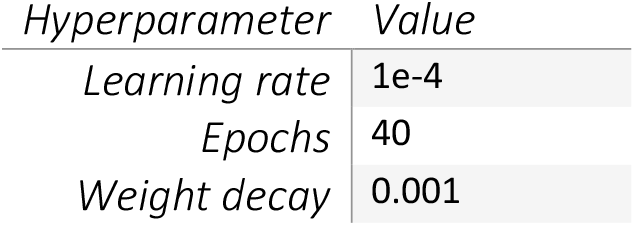
Hyperparameters for all models for Method 1.

### Method 2

**Table S2.** Hyperparameter values for Ankh large with weighted cross entropy loss.

| Hyperparameter | Value |
| --- | --- |
| rank | 4 |
| Lora alpha | 1 |
| Bias | None |
| Lora dropout | 0.1 |
| LoRA Layers | key, query, value |
| Learning rate | 3e-4 |
| Weight decay | 0.2 |
| LR scheduler | Cosine |
| Epochs | 30 |

**Table S3.** Hyperparameter values for Ankh large with focal loss.

| Hyperparameter | Value |
| --- | --- |
| rank | 8 |
| Lora alpha | 16 |
| Bias | None |
| Lora dropout | 0.1 |
| LoRA Layers | key, query, value, output |
| Learning rate | 3e-4 |
| Weight decay | 0.2 |
| LR scheduler | Cosine |
| Epochs | 30 |

**Table S4.** Hyperparameter values ESM-2 3B with weighted cross entropy loss.

| Hyperparameter | Value |
| --- | --- |
| *rank* | 8 |
| *Lora alpha* | 16 |
| *Bias* | None |
| *Lora dropout* | 0.1 |
| *LoRA Layers* | key, query, value |
| *Learning rate* | 3e-4 |
| *Weight decay* | 0.2 |
| *LR scheduler* | Cosine |
| *Epochs* | 30 |

**Table S5.** Hyperparameter values ESM-2 3B with focal loss.

| *Hyperparameter* | *Value* |
| --- | --- |
| *rank* | 8 |
| *Lora alpha* | 16 |
| *Bias* | None |
| *Lora dropout* | 0.0 |
| *LoRA Layers* | key, query, value, dense |
| *Learning rate* | 3e-4 |
| *Weight decay* | 0.2 |
| *LR scheduler* | Cosine |
| *Epochs* | 30 |

**Table S6.** Hyperparameter values for ProtT5 XL for both weighted cross entropy and focal loss.

| *Hyperparameter* | *Value* |
| --- | --- |
| *rank* | 8 |
| *Lora alpha* | 16 |
| *Bias* | None |
| *Lora dropout* | 0.1 |
| *LoRA Layers* | key, query, value |
| *Learning rate* | 3e-4 |
| *Weight decay* | 0.2 |
| *LR scheduler* | Cosine |
| *Epochs* | 30 |

### Method 3

**Table S7.** Hyperparameter values for Ankh large.

| *Hyperparameter* | *Value* |
| --- | --- |
| *rank* | 4 |
| *Lora alpha* | 1 |
| *Bias* | None |
| *Lora dropout* | 0.5 |
| *LoRA Layers* | key, query, value |
| *Learning rate* | 3e-4 |
| *Weight decay* | 0.5 |
| *LR scheduler* | Cosine |
| *Epochs* | 5 |

**Table S8.** Hyperparameter values ESM-2 3B.

| *Hyperparameter* | *Value* |
| --- | --- |
| *rank* | 8 |
| *Lora alpha* | 16 |
| *Bias* | None |
| *Lora dropout* | 0.25 |
| *LoRA Layers* | key, query, value |
| *Learning rate* | 3e-4 |
| *Weight decay* | 0.4 |
| *LR scheduler* | Cosine |
| *Epochs* | 5 |

**Table S9.** Hyperparameter values for ProtT5 XL.

| *Hyperparameter* | *Value* |
| --- | --- |
| *rank* | 8 |
| *Lora alpha* | 16 |
| *Bias* | None |
| *Lora dropout* | 0.25 |
| *LoRA Layers* | key, query, value |
| *Learning rate* | 3e-4 |
| *Weight decay* | 0.4 |
| *LR scheduler* | Cosine |
| *Epochs* | 10 |

## References

[1] J. Guo and H.-X. Zhou, “Protein Allostery and Conformational Dynamics.,” Chemical ReviewS Journal, vol. 116, no. 11, pp. 6503–6515, 2016.

[2] L. G. Ahuja, S. S. Taylor and A. P. Kornev, “Tuning the “violin” of protein kinases: The role of dynamics-based allostery,” IUBMB Life, vol. 71, no. 6, pp. 685–696, 2019.

[3] M. B. N. R. Gunasekaran K, “Is allostery an intrinsic property of all dynamic proteins?,” Proteins, vol. 57, no. 3, pp. 433–443, 2004.

[4] T. C. Nussinov R, “The different ways through which specificity works in orthosteric and allosteric drugs,” Curr Pharm Des, vol. 18, no. 0, pp. 1311–6, 2012.

[5] K. F. Rudolph U, “Beyond classical benzodiazepines: novel therapeutic potential of GABAA receptor subtypes,” Nat Rev Drug Discov, vol. 10, no. 9, pp. 685–97, 2011.

[6] Z. Huang, L. Zhu, Y. Cao, G. Wu, X. Liu, Y. Chen, Q. Wang, T. Shi, Y. Zhao, Y. Wang, W. Li, Y. Li, H. Chen, G. Chen and J. Zhang, “ASD: a comprehensive database of allosteric proteins and modulators.,” Nucleic Acids Research, vol. 39, pp. 663–669, 2011.

[7] C. Z. Nerín-Fonz F, “Machine learning approaches in predicting allosteric sites,” Current Opinion in Structural Biology, vol. 85, p. 102774, 2024.

[8] V. Le Guilloux, P. Schmidtke and P. Tuffery, “Fpocket: An open source platform for ligand pocket detection,” BMC Bioinformatics, vol. 10, no. 168, 2009.

[9] Z. Huang, L. Mou, Q. Shen, S. Lu, C. Li, X. Liu, G. Wang, S. Li, L. Geng, Y. Liu, J. Wu, G. Chen and J. Zhang, “ASD v2.0: updated content and novel features focusing on allosteric regulation.,” Nucleic Acids Research, vol. 42, pp. 510–516, 2014.

[10] W. Huang, G. Wang, Q. Shen, X. Liu, S. Lu, L. Geng, Z. Huang and J. Zhang, “ASBench: benchmarking sets for allosteric discovery,” Bioinformatics, vol. 31, no. 15, pp. 2598–6000, 2015.

[11] Q. Shen, G. Wang, S. Li, X. Liu, S. Lu, Z. Chen, K. Song, J. Yan, L. Geng, Z. Huang, W. Huang, G. Chen and J. Zhang, “ASD v3.0: unraveling allosteric regulation with structural mechanisms and biological networks,” Nucleic Acids Research, vol. 44, no. D1, p. D527–D535, 2016.

[12] X. Liu, S. Lu, K. Song, Q. Shen, D. Ni, Q. Li, X. He, H. Zhang, Q. Wang, Y. Chen, X. Li, J. Wu, C. Sheng, G. Chen, Y. Liu, X. Lu and J. Zhang, “Unraveling allosteric landscapes of allosterome with ASD,” Nucleic Acids Research, vol. 48, no. D1, p. D394–D401, 2020.

[13] W. Huang, S. Lu, Z. Huang, X. Liu, L. Mou, Y. Luo, Y. Zhao, Y. Liu, Z. Chen, T. Hou and J. Zhang, “Allosite: a method for predicting allosteric sites,” Bioinformatics, vol. 29, no. 18, pp. 2357–2359, 2013.

[14] J. Greener and M. Sternberg, “AlloPred: prediction of allosteric pockets on proteins using normal mode perturbation analysis,” BMC Bioinformatics, vol. 16, no. 335, 2015.

[15] K. Song, X. Liu, W. Huang, S. Lu, Q. Shen, L. Zhang and Z. J., “Improved Method for the Identification and Validation of Allosteric Sites,” Journal of Chemical Information and Modeling, vol. 57, no. 9, pp. 2358–2363, 2017.

[16] R. Akbar and V. Helms, “ALLO: A tool to discriminate and prioritize allosteric pockets,” Chemical Biology and Drug Design, vol. 91, no. 4, pp. 845–853, 2018.

[17] H. Tian, S. Xiao, X. Jiang and P. Tao, “PASSer: fast and accurate prediction of protein allosteric sites,” Nucleic Acids Research, vol. 51, no. W1, p. W427–W431, 2023.

[18] S. Xiao, H. Tian and P. Tao, “PASSer2.0: Accurate Prediction of Protein Allosteric Sites Through Automated Machine Learning,” Frontiers in Molecular Biosciences, vol. 9, 2022.

[19] H. Tian, S. Xiao, X. Jiang and T. P, “PASSerRank: Prediction of Allosteric Sites with Learning to Rank,” ArXiv [Preprint], vol. 44, no. 28, pp. 2223–2229, 2023.

[20] J. Zha, Q. Li, X. Liu, W. Lin, T. Wang, J. Wei, Z. Zhang, X. Lu, J. Wu, D. Ni, K. Song, L. Zhang, X. Lu, S. Lu and J. Zhang, “AlloReverse: multiscale understanding among hierarchical allosteric regulations,” Nucleic Acids Research, vol. 51, no. W1, p. W33–W38, 2023.

[21] W. Yan, G. L. Z. Hu, J. Zhou, Y. Yang and J. S. B. Chen, “Node-Weighted Amino Acid Network Strategy for Characterization and Identification of Protein Functional Residues,” Journal of Chemical Information and Modeling, vol. 58, no. 9, pp. 2024–2032, 2018.

[22] S. Mishra, G. Kandoi and R. Jernigan, “Coupling dynamics and evolutionary information with structure to identify protein regulatory and functional binding sites,” PROTEINS: Structure, Function, and Bioinformatics, vol. 87, pp. 850–868, 2019.

[23] J. Xie, G. Pan, Y. Li and L. Lai, “How protein topology controls allosteric regulations,” Journal of Chemical Physics, vol. 158, no. 10, p. 105102, 2023.

[24] A. Elnaggar, H. Essam, W. Salah-Eldin, W. Moustafa, M. Elkerdawy, C. Rochereau and B. Rost, “Ankh: Optimized Protein Language Model Unlocks General-Purpose Modelling,” arXiv, 2023.

[25] Z. Lin, H. Akin, R. Rao, B. Hie, Z. Zhu, W. Lu, N. Smetanin, R. Verkuil, O. Kabeli, Y. Shmueli, A. Dos Santos Costa, M. Fazel-Zarandi, S. Candido and A. Rives, “Evolutionary-scale prediction of atomic-level protein structure with a language model,” Science, vol. 379, no. 6637, pp. 1123–1130, 2023.

[26] N. Brandes, D. Ofer, Y. Peleg, N. Rappoport and M. Linial, “ProteinBERT: a universal deep-learning model of protein sequence and function,” Bioinformatics, vol. 38, no. 8, p. 2102–2110, 2022.

[27] P. P. Ray, “ChatGPT: A comprehensive review on background, applications, key challenges, bias, ethics, limitations and future scope,” Internet of Things and Cyber-Physical Systems, vol. 3, pp. 121–154, 2023.

[28] A. Vaswani, N. Shazeer, N. Parmar, K. Uszkoreit, L. Jones, A. Gomez, L. Kaiser and I. Polosukhin, “Attention Is All You Need,” Advances in Neural Information Processing Systems 30 (NIPS 2017), 2017.

[29] J. Devlin, M.-W. Chang, K. Lee and K. Toutanova, “BERT: Pre-training of Deep Bidirectional Transformers for Language Understanding,” ArXiv, 2018.

[30] D. Ofer, N. Brandes and M. Linial, “The language of proteins: NLP, machine learning & protein sequences,” Computational and Structural Biotechnology Journal, vol. 19, pp. 1750–1758, 2021.

[31] Z. Liu, Y. Li, L. Han, J. Li, J. Liu, Z. Zhao, W. Nie, Y. Liu and R. Wang, “PDB-wide collection of binding data: current status of the PDBbind database,” Bioinformatics, vol. 31, no. 3, pp. 405–412, 2015.

[32] Z. Liu, M. Su, L. Han, J. Liu, Q. Yang, Y. Li and R. Wang, “Forging the Basis for Developing Protein-Ligand Interaction Scoring Functions,” Accounts of Chemical Research, vol. 50, no. 2, pp. 302–309, 2017.

[33] D. A. Bloore, J. Kim, K. Kapoor, E. Chen, K. Gao, M. Wang and M. Hao, “Protein Language Models Enable Accurate Cryptic Ligand Binding Pocket Prediction,” in The NeurIPS 2023 Workshop on New Frontiers of AI for Drug Discovery and Development (AI4D3 2023), New Orleans, LA, USA, 2023.

[34] P. Cimermancic, P. Weinkam, T. Rettenmaier, L. Bichmann, D. Keedy, R. Woldeyes, D. Schneidman-Duhovny, O. Demerdash, J. Mitchell, J. Wells, J. Fraser and A. Sali, “CryptoSite: Expanding the Druggable Proteome by Characterization and Prediction of Cryptic Binding Sites.,” J Mol Biol., vol. 428, no. 4, pp. 709–719, 2016.

[35] C. Raffel, N. Shazeer, A. Roberts, K. Lee, S. Narang, M. Matena, Y. Zhou, W. Li and P. J. Liu, “Exploring the Limits of Transfer Learning with a Unified Text-to-Text Transformer,” Journal of Machine Learning Research, vol. 21, no. 140, pp. 1–67, 2020.

[36] M. Steinegger and J. Söding, “MMseqs2 enables sensitive protein sequence searching for the analysis of massive data sets,” Nature Biotechnology, vol. 35, p. 1026–1028, 2017.

[37] Z. Liu, M. Su, L. Han, J. Liu, Q. Yang, Y. Li and R. Wang, “Forging the Basis for Developing Protein-Ligand Interaction Scoring Functions,” Accounts of Chemical Research, vol. 50, no. 2, pp. 302–309, 2017.

[38] M. Hauser, M. Steinegger and J. Söding, “MMseqs software suite for fast and deep clustering and searching of large protein sequence sets.,” Bioinformatics, vol. 32, no. 9, pp. 1323–1330, 2016.

